## Supplemental Material for "POT1 Stability and Binding Measured by Fluorescence Thermal Shift Assays"

### Captions

**Scheme S1.** Equations used for the simulation and analysis of FTSA data.

**Figure S1.** The FTSA assay can be used for the quality control assessment of POT1 preparations. The data show 22 preparations of POT1 alone (A) or in the presence of oligonucleotide O1 (B). Binding of oligonucleotide O1 defines a function of POT1. It is clear that 4 of the 22 preparations show anomalous binding. These preparations were not used in any subsequent experimental studies. The origin of the altered function is not known. For all curves, [POT1] = 5 μM, [O1] = 50 μM.

**Figure S2.** Stability of POT1 in the presence of a variety of cosolutes. The difference in the transition melting temperature is shown with reference to 51.5 $^{o}$C.

**Figure S3**. Determination of POT1 thermal denaturation thermodynamics. Data were obtained by FTSA (A) or independently by circular dichroism (B). (A) Primary FTSA data were fit to a two-state denaturation model that included parameters for pre- and post- transition baselines. A global fit was done for 8 experiments, linking $\Delta H$ and $T_m$ for all data sets while allowing the baseline parameters to adjust for each individual experiment. The residual plots of the fits are shown as the curves centered at Y= 0. (B) Derivative denaturation curves for two experiments obtained by circular dichroism. Data were fit using the two-state method of John and Weeks.[1]

**Figure S4.** Congo Red and putative POT1 binding compounds predicted by virtual screening.

**Figure S5**. Thermal shift data for putative POT1 binding compounds. (A). $\Delta T_m$ values for 91 compounds predicted by virtual screening to bind to POT1. (B) Distribution of $\Delta T_m$ values shown as a box plot. The mean is shown as the black square. The top and bottom edges of the square are the 75$^{th}$ and 25$^{th}$ percentiles. The open circles show the 1$^{st}$ and 99$^{th}$ percentiles.

**Figure S6**. Nucleotide analogues tested in POT1 thermal denaturation studies. (A) Table of analogues tested with corresponding $\Delta T_m$ values derived from POT1 melting curves in B. (B) Representative thermal denaturation curves of POT1 in the presence of DMSO (control) or nucleotides shown in A.

**Figure S7**. Analysis of POT1-O1 binding isotherm using the simple bimolecular binding model proposed in reference[2].

### Scheme S1

$$U + L_f \underset{Folding}{\overset{K_U}{\rightleftharpoons}} N + L_f \underset{Binding}{\overset{K_b}{\rightleftharpoons}} NL_b$$

$$[\mathrm{L}]_t = (K_U - 1)\left(\frac{1}{K_b} + \frac{[\mathrm{P}]_t}{2K_U}\right)$$

$$K_U = \exp\left(-\frac{\Delta H^{\circ}_{U\_T_r} + \Delta C_{p\_U}(T_m - T_r) - T_m\left(\Delta S^{\circ}_{U\_T_r} + \Delta C_{p\_U}\ln\frac{T_m}{T_r}\right)}{RT_m}\right)$$

$$K_b = \exp\left(-\frac{\Delta H^{\circ}_{b\_T_0} + \Delta C_{p\_b}(T_m - T_0) - T_m\left(\Delta S^{\circ}_{b\_T_0} + \Delta C_{p\_b}\ln\frac{T_m}{T_0}\right)}{RT_m}\right)$$

### Definitions of Symbols

| Parameter | Description | Units |
|---|---|---|
| $[\mathrm{P}]_t$ | Total protein concentration | M |
| $[\mathrm{L}]_t$ | Total ligand concentration | M |
| $K_U$ | Equilibrium constant, unfolding | *Dimensionless* |
| $K_b$ | Equilibrium constant, binding | $M^{-1}$ |
| $T_m$ | Temperature at transition midpoint in presence of ligand | K |
| $T_r$ | Temperature at transition midpoint of protein denaturation | K |
| $T_0$ | Reference temperature of ligand binding parameters | K |
| $\Delta H^{\circ}_{U\_T_r}$ | Enthalpy of protein denaturation at the transition midpoint temperature | J $mol^{-1}$ |
| $\Delta C_{p\_U}$ | Heat capacity change for protein denaturation | J $mol^{-1}K^{-1}$ |
| $\Delta S^{\circ}_{U\_T_r}$ | Entropy of protein denaturation at the transition midpoint temperature | J $mol^{-1}K^{-1}$ |
| $\Delta H^{\circ}_{b\_T_0}$ | Enthalpy of ligand binding at $T_0$ | J $mol^{-1}$ |
| $\Delta C_{p\_b}$ | Heat capacity change for ligand binding | J $mol^{-1}K^{-1}$ |
| $\Delta S^{\circ}_{b\_T_0}$ | Entropy of ligand binding at $T_0$ | J $mol^{-1}K^{-1}$ |
| R | Universal gas constant (8.3144598) | J $mol^{-1}K^{-1}$ |

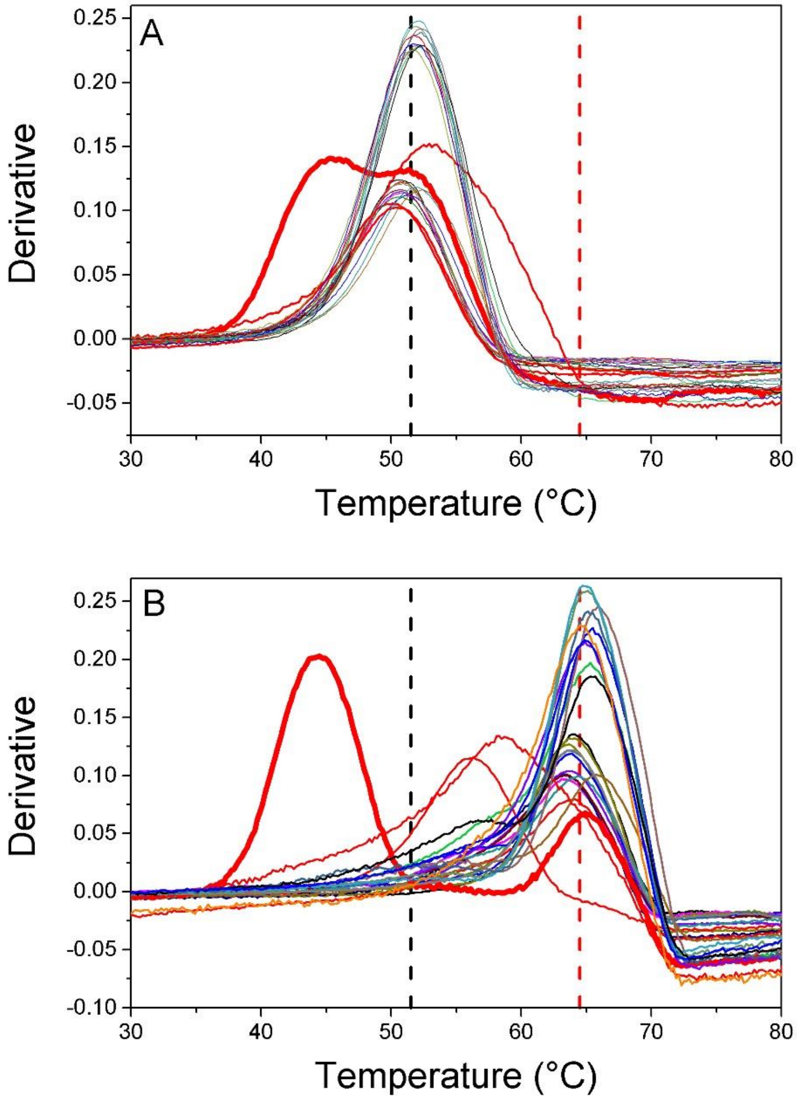


**Figure S1.** The DSF assay can be used for the quality control assessment of POT1 preparations. The data show 22 preparations of POT1 alone (A) or in the presence of oligonucleotide O1 (B). Binding of oligonucleotide O1 defines a function of POT1. It is clear that 4 of the 22 preparations show anomalous binding. These preparations were not used in any subsequent experimental studies. The origin of the altered function is not known. For all curves, [POT1] = 5 μM, [O1] = 50 μM.

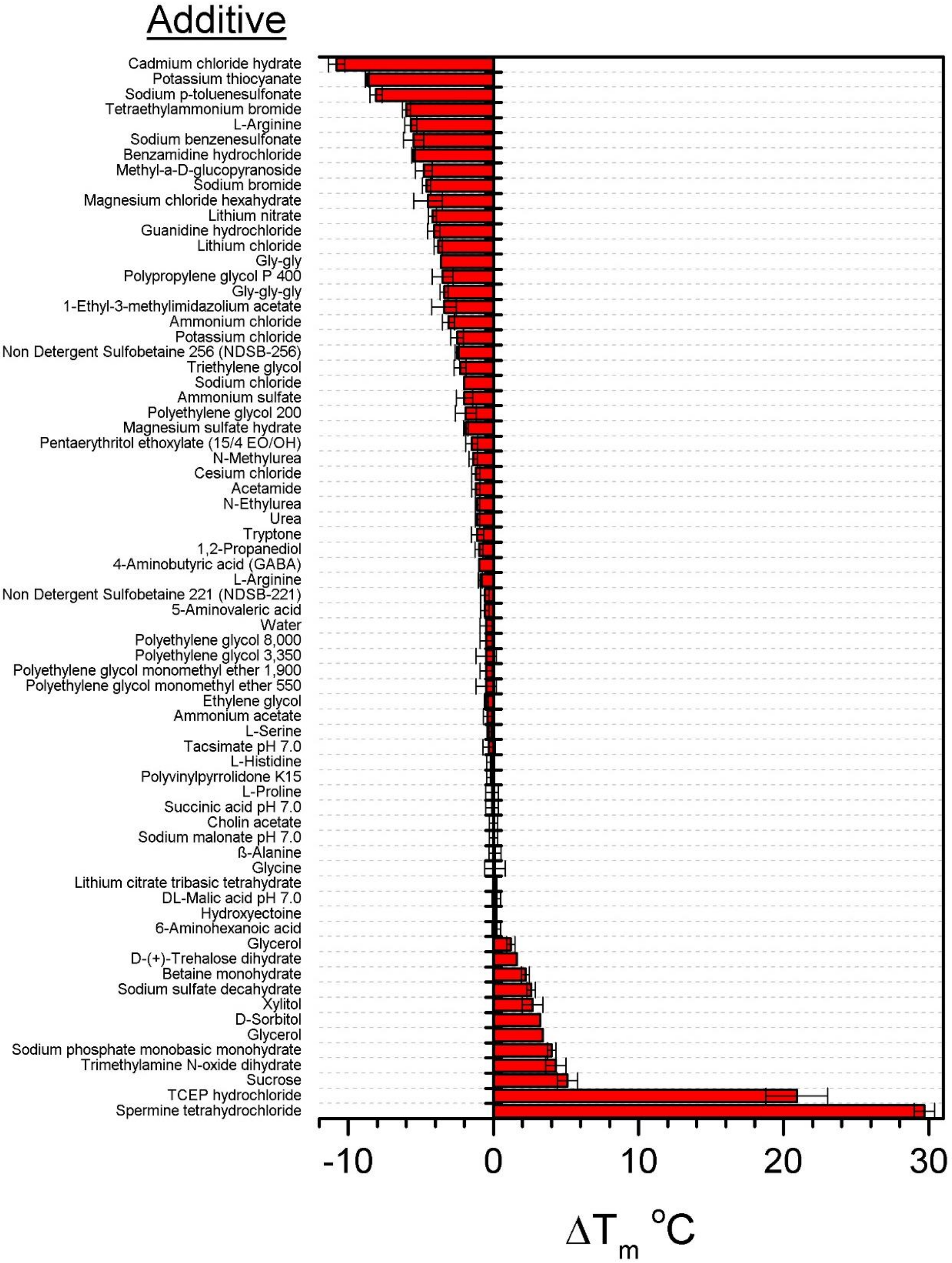


**Figure S2.** Stability of POT1 in the presence of a variety of cosolutes. The difference in the transition melting temperature is shown with reference to 51.5 °C. These results were obtained using the Solubility & Stability Screen kit from Hampton Research Corp. (Viejo, CA). Full details of the reagents and concentrations of additives can be found at: https://hamptonresearch.com/uploads/support_materials/HR2-072_Binder.pdf.

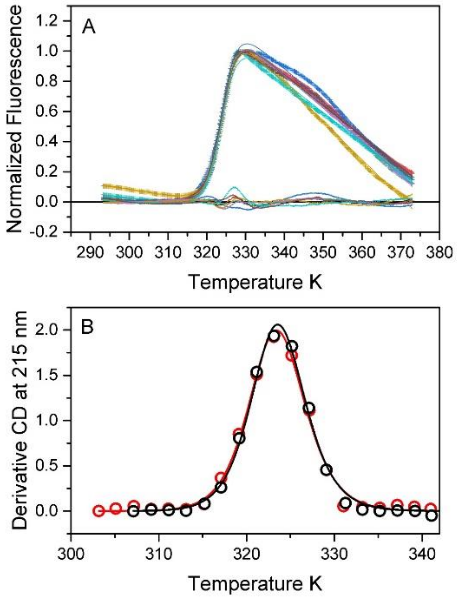


**Figure S3**. Determination of POT1 thermal denaturation thermodyamics. Data were obtained by FTSA (A) or independently by circular dichroism (B). (A) Primary FTSA data were fit to a two-state denaturation model that included parameters for pre- and post- transition baselines. A global fit was done for 8 experiments, linking $\Delta H$ and $T_m$ for all data sets while allowing the baseline parameters to adjust for each individual experiment. The residual plots of the fits are shown as the curves centered at Y= 0. (B) Derivative denaturation curves for two experiments obtained by circular dichroism. Data were fit using the two-state method of John and Weeks[1].

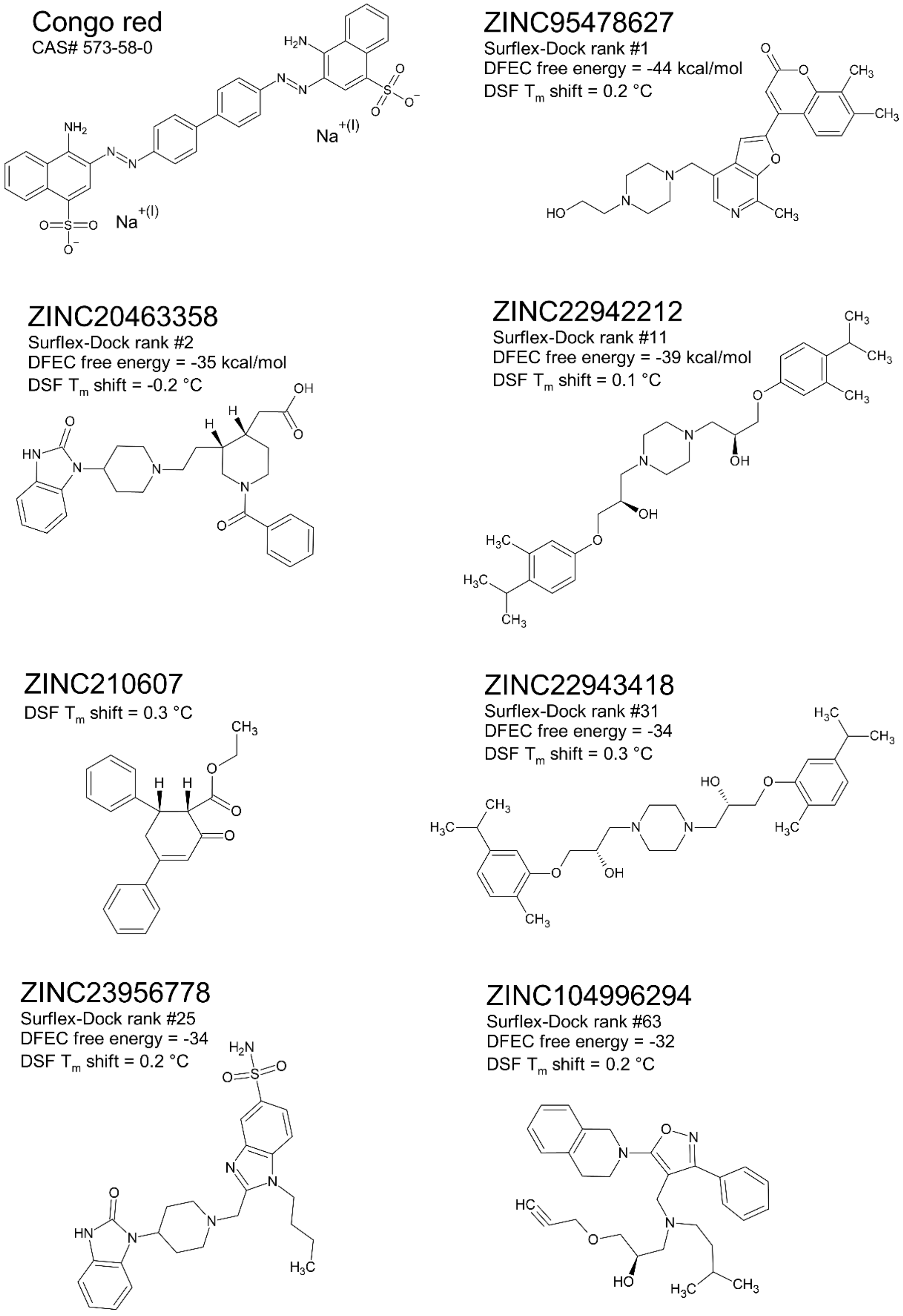


**Figure S4.** Congo Red and representative POT1 binding compounds predicted by virtual screening.

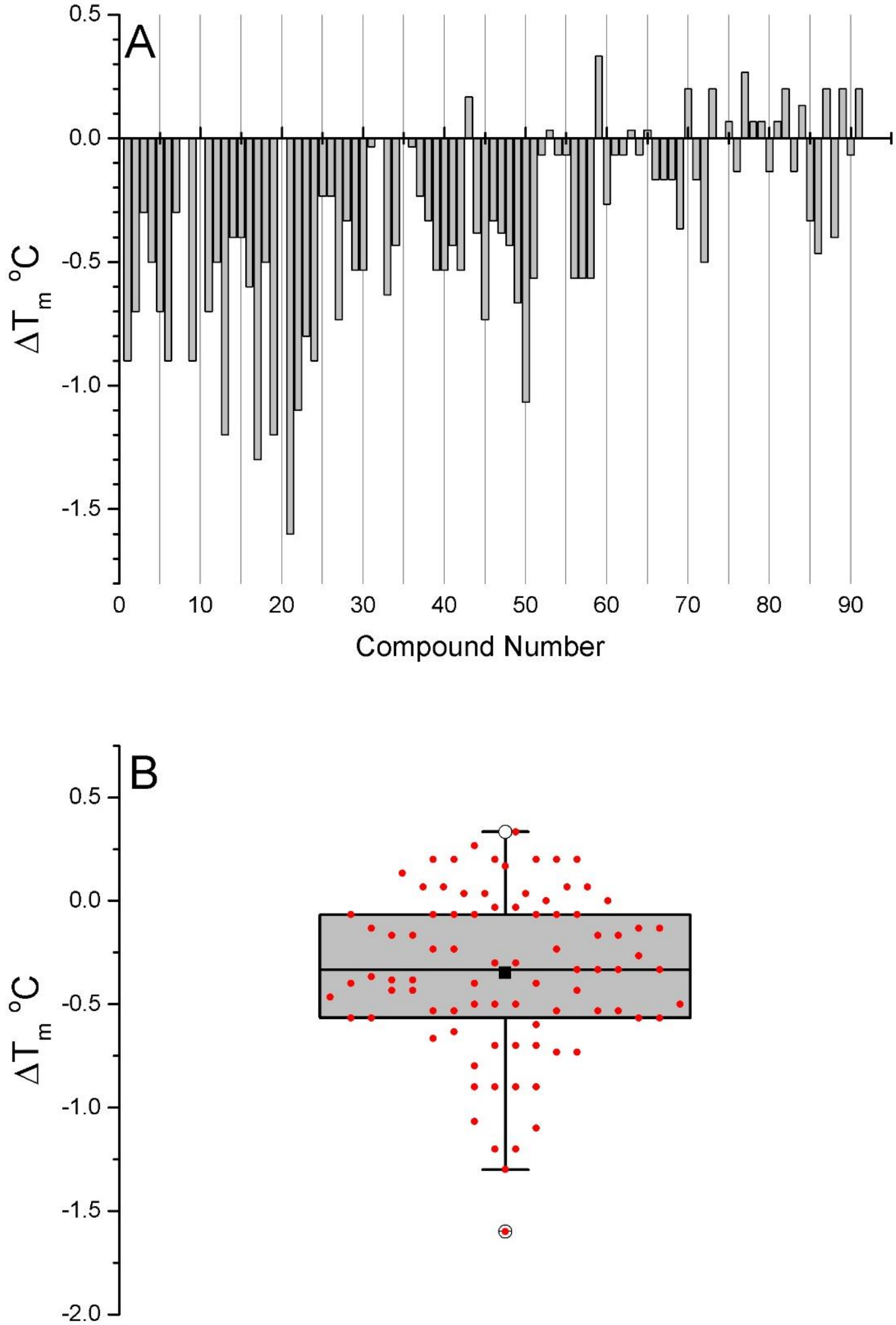


**Figure S5**. Thermal shift data for putative POT1 binding compounds. (A). $\Delta T_m$ values for 91 compounds predicted by virtual screening to bind to POT1. (B) Distribution of $\Delta T_m$ values shown as a box plot. The mean is shown as the black square. The top and bottom edges of the square are the 75th and 25th percentiles. The open circles show the 1st and 99th percentiles.

A

| Compound | Structure | $\Delta T_m$ (°C) | Compound | Structure | $\Delta T_m$ (°C) |
|---|---|---|---|---|---|
| Decitabine | 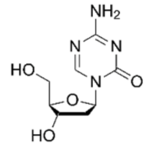 | 1.0 | Fludarabine | 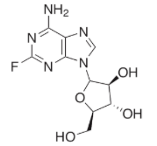 | 1.4 |
| Gemcitabine | 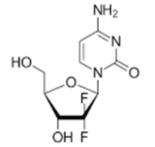 | 1.6 | Capecitabine | 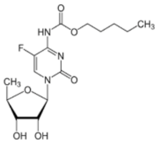 | 1.6 |
| Fluorouracil | 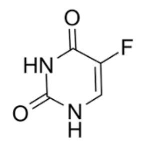 | 1.2 | Azacitidine | 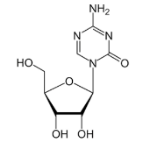 | 1.8 |
| Cladribine | 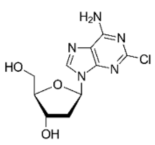 | 1.2 | Pentostatin | 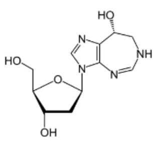 | 1.4 |
| Clofarabine | 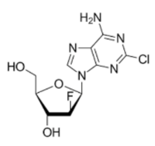 | 1.6 | Nelarabine | 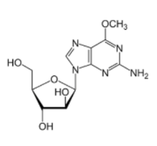 | 1.8 |

B

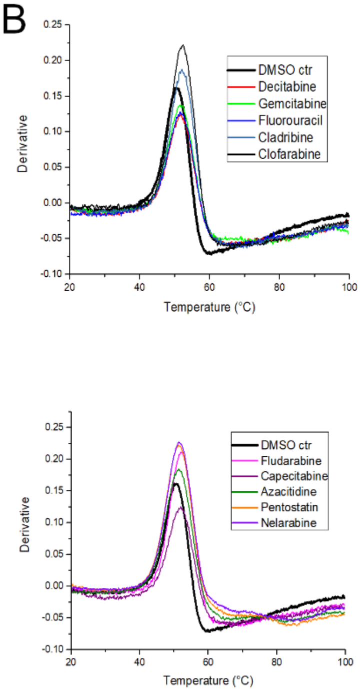


**Figure S6**. Nucleotide analogues tested in POT1 thermal denaturation studies. (A) Table of analogues tested with corresponding $\Delta T_m$ values derived from POT1 melting curves in B. (B) Representative thermal denaturation curves of POT1 in the presence of DMSO (control) or nucleotides shown in A.

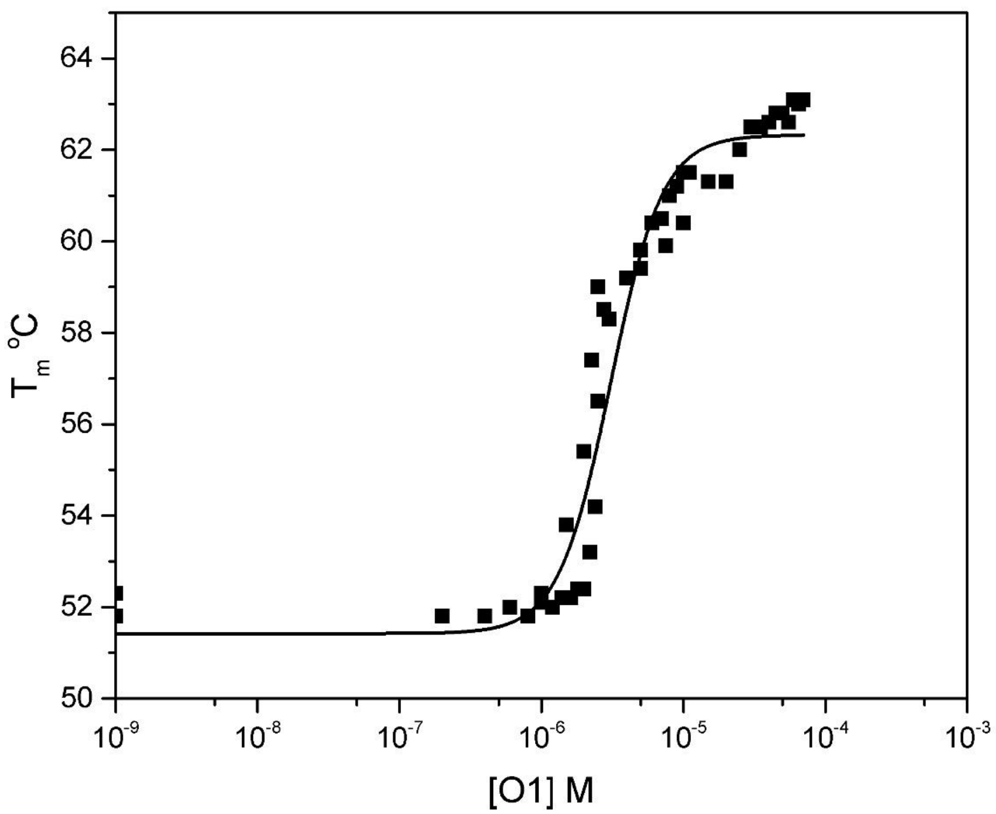


**Figure S7**. Analysis of POT1-O1 binding isotherm using the simple bimolecular binding model proposed by Vivoli et al [2].

**Table S1**. Thermodynamics of POT1 thermal denaturation.

| **Method** | **$T_m$<br>ºK** | **$\Delta H_{vH}$<br>kJ $mol^{-1}$** | **$\Delta S$<br>J $mol^{-1}K^{-1}$** | **Comments** |
|---|---|---|---|---|
| FTSA | 324.06 ± 0.01<br><324.05;324.08> | 442 ± 2<br><4440; 444> | 1364 ± 7 | Global fit to 8 FTSA experiments |
| CD<br>A<br>B | 323.4±0.1<br>323.5±0.1<br><323.2;323.8> | 410 ± 11<br>425 ± 18<br><386; 462> | 1266<br>1311 | Single wavelength: 215 nm |
| CD | 323.9 | 425 | 1315 | Fit to amplitude vector from SVD analysis of spectra collected from 200-280 nm |

The 65% confidence interval is indicated by the brackets <LCL; UCL>

**Table S2.** Oligonucleotides used in this study.

| Name | Sequence* | Structure in KCl | $\varepsilon$260 nm ($mM^{-1}$ $cm^{-1}$) | nt |
|---|---|---|---|---|
| 143D | d[A(GGGTTA)$_3$GGG] | hybrid-1+2 | 228.5 | 22 |
| 2GKU | d[TT(GGGTTA)$_3$GGGA] | hybrid-1 | 244.3 | 24 |
| 2JSL | d[TA(GGGTTA)$_3$GGG]TT | hybrid-2 | 261.2 | 25 |
| 1XAV | d[TGAGGGTGGGTAGGGTGGGTAA] | parallel | 228.7 | 22 |
| O1 | d[TTAGGGTTAG] | single-strand | 102.2 | 10 |
| O1c | d[CTAACCCTAA] | single-strand | 96.6 | 10 |
| PNA | TTAGGGTTAG-O | single-strand | 109.4 | 10 |

*FRET-oligo contains 5’-6FAM and 3’-TAMRA labels

**Table S3.** Variations of O1 sequences

| O1 Variations | Sequence |
|---|---|
| MBS+1 | d[TTA GGG TTA G] |
| MBS | d[TAG GGT TAG] |
| MBS-T7A | d[TAG GGT AAG] |
| MBS-A8T | d[TAG GGT TTG] |
| MBS-G5C | d[TAG GCT TAG] |
| MBS-T6A | d[TAG GGA TAG] |
| Permut4 | d[GTT AGG GTT] |
| Permut3 | d[GGT TAG GGT] |
| MBS-G4C | d[TAG CGT TAG] |
| Permut5 | d[TTA GGG TTA] |
| MBS-A2T | d[TTG GGT TAG] |
| MBS-G9C | d[TAG GGT TAC] |
| MBS-G3C | d[TAC GGT TAG] |
| MBS-G9A | d[TAG GGT TAA] |
| MBS-T1A | d[CAG GGT TAG] |
| Permut1 | d[AGG GTT AGG] |
| Permut2 | d[GGG TTA GGG] |
| MBS-G9C' | d[TAG GGT TAC] |
